# Temporal Multivariate Pattern Analysis (tMVPA): a single trial approach exploring the temporal dynamics of the BOLD signal

**DOI:** 10.1101/273110

**Authors:** Luca Vizioli, Alexander Bratch, Junpeng Lao, Kamil Ugurbil, Lars Muckli, Essa Yacoub

## Abstract

**Background:** fMRI provides spatial resolution that is unmatched by any non-invasive neuroimaging technique. Its temporal dynamics however are typically neglected due to the sluggishness of the hemodynamic based fMRI signal.

**New Methods:** We present *temporal multivariate pattern analysis* (tMVPA), a method for investigating the temporal evolution of neural representations in fMRI data, computed using pairs of single-trial BOLD time-courses, leveraging both spatial and temporal components of the fMRI signal. We implemented an expanding sliding window approach that allows identifying the time-window of an effect.

**Results:** We demonstrate that tMVPA can successfully detect condition-specific multivariate modulations over time, in the absence of univariate differences. Using Monte Carlo simulations and synthetic data, we quantified family-wise error rate (FWER) and statistical power. Both at the group and at the single subject level, FWER was either at or significantly below 5%. For the group level, we reached the desired power with 18 subjects and 12 trials; for the single subject scenario, 14 trials were required to achieve comparable power.

**Comparison with existing methods:** tMVPA adds a temporal multivariate dimension to the tools available for fMRI analysis, enabling investigations of the evolution of neural representations over time. Moreover, tMVPA permits performing single subject inferential statistics by considering single-trial distribution.

**Conclusion:** The growing interest in fMRI temporal dynamics, motivated by recent evidence suggesting that the BOLD signal carries temporal information at a finer scale than previously thought, advocates the need for analytical tools, such as the tMVPA approach proposed here, tailored to investigating BOLD temporal information.

## Introduction

Over the past quarter century, functional Magnetic Resonance Imaging (fMRI) has become one of the most powerful non-invasive tools for investigating human neural processing. By exploiting the coupling between oxygenated blood flow and neuronal firing (Goense & Logothetis, 2008; Logothetis, N. K., Pauls, J., Augath, M., Trinath, T., & Oeltermann, 2001; S Ogawa et al., 1993), fMRI infers cortical activity by measuring changes in the Blood Oxygen Level Dependent (BOLD) signal (Goense & Logothetis, 2008; Logothetis, N. K., Pauls, J., Augath, M., Trinath, T., & Oeltermann, 2001; S Ogawa et al., 1993). The sluggish nature of the hemodynamic based BOLD signal (requiring several seconds to peak following stimulus presentation(Boynton et al., 1996; S Ogawa et al., 1993)), paired with the high spatial precision of fMRI recordings, has resulted in a focus on BOLD spatial information in most applications, neglecting any temporal dynamics. More recently, developments in fMRI pulse sequences, allowing significant increases in temporal resolution (Feinberg, D. A., Moeller, S., Smith, S. M., Auerbach, E., Ramanna, S., Glasser, M. F., … & Yacoub, 2010; Moeller et al., 2010) that have been thus far primarily exploited to improve statistical power in fMRI analysis, offer the possibility of resolving temporal dynamics that were previously elusive.

While focus has been primarily on the spatial domain of the BOLD signal, this is not to say that the fMRI temporal domain has been entirely ignored. For example, several attempts have been made to target local stimulus-distinct characteristics of the BOLD time series. Specifically, these investigations have sought to understand stimulus-specific temporal effects in the context of decision making (Mcguire & Kable, 2015), auditory (Baumann et al., 2010), and semantic and visual processing (Avossa et al., 2003; Bailey et al., 2013; Formisano et al., 2002; Gentile et al., 2017; Siero, J.C., Petridou, N., Hoogduin, H., Luijten, P.R., Ramsey, 2011; Vu, A.T., Phillips, J.S., Kay, K., Phillips, M.E., Johnson, M.R., Shinkareva, S.V., Tubridy, S., Millin, R., Grossman, M., Gureckis, T., Bhattacharyya, R., Yacoub, 2016). In conjunction, animal studies have sought to understand the precise relationship between the BOLD temporal dynamics and the neural activity elicited from such domains (Silva & Koretsky, 2002; Yen et al., 2018). Additionally, it is worth noting that a variety of both high complexity and real world stimuli operate at the temporal resolution available to fMRI. For example, in the visual domain, a number of visual illusions are characterized by their slowly transforming, bi-stable nature (Ernst & Bu, 2004; Schrater et al., 2004). Furthermore, biological motion (Johansson, 1973; Maier et al., 2008; Troje, 2002) and other motion-based complex stimuli (Ball & Sekuler, 1982; Shadlen & Newsome, 1998) are typically presented over large temporal windows. BOLD latency measurements have likewise been shown to be relevant in the auditory and multisensory domain, where, for example, phonemic boundaries shift across temporal gradients when presented in isolation (Lee et al., 2012) or within specific visual contexts (Gribble, 1996). Moreover, analyses of neural responses to any long duration stimuli, such as film or real-world dynamic scenes, necessitate a technique that directly measures the temporal evolution of the BOLD signal.

Importantly, a number of studies have more recently suggested that fMRI may carry neuronal information at a much faster temporal scale than previously (Lewis et al., 2016; Siero, J.C., Petridou, N., Hoogduin, H., Luijten, P.R., Ramsey, 2011; Vu, A.T., Phillips, J.S., Kay, K., Phillips, M.E., Johnson, M.R., Shinkareva, S.V., Tubridy, S., Millin, R., Grossman, M., Gureckis, T., Bhattacharyya, R., Yacoub, 2016). Siero and colleagues (Siero, J.C., Petridou, N., Hoogduin, H., Luijten, P.R., Ramsey, 2011), for example, indicated that neurovascular coupling takes place on a shorter timescale than had been previously reported in the human brain. Moreover, Lewis and colleagues (Lewis et al., 2016) have suggested that, due to recent advances in MR hardware and software as well as analytical strategies, fMRI can measure neural oscillations up to 1 Hz. Additionally, Vu and colleagues (Vu, A.T., Phillips, J.S., Kay, K., Phillips, M.E., Johnson, M.R., Shinkareva, S.V., Tubridy, S., Millin, R., Grossman, M., Gureckis, T., Bhattacharyya, R., Yacoub, 2016) successfully demonstrated that with the use of multivoxel pattern analysis (MVPA), it is possible to extract word timing information with fast TRs (i.e. 500 ms). Along the same lines, in an visual illusion experiment, Edwards and colleagues (Edwards et al., 2017) showed that as little as 32 ms difference in stimulus presentation is reliably detected in the BOLD time-course.

These observations highlight the growing interest in the temporal dynamics of the BOLD signal. However, to fully exploit the potential neuro-temporal information carried by the BOLD time-course, MR hardware and software (e.g. pulse sequences) developments have to be paired with suitable analytical tools that maximize the sensitivity to BOLD temporal information. Thus far, the majority of temporal analyses have only examined univariate temporal differences between stimuli or stimulus conditions (i.e., latency differences on average amplitude). While such data is useful for understanding the propagation of neural activation throughout the brain as a function of time, it fails to capture the representational content as conveyed by multivariate patterns as well as how these representations transform over time. Multivariate approaches to analyzing fMRI data offer a different, albeit complementary outlook on the neural information carried by the BOLD signal (Kriegeskorte & Bandettini, 2007). It has been suggested that multivoxel pattern analysis, or MVPA (Haxby et al., 2001; Kamitani & Tong, 2005), has the ability to optimally probe neuronal information existing in voxel populations with conventional fMRI methods (Carlson et al., 1999; Cox & Savoy, 2003; Haxby et al., 2005; Kriegeskorte & Bandettini, 2007; Strother et al., 2002). Even at 3T, where voxels traditionally measure 2-3 mm isotropic resolutions, MVPA can successfully extract neural information – such as orientation preference (Kamitani & Tong, 2005) – which exists at a much finer spatial scale than the resolution of single voxels. These approaches are believed to increase the sensitivity to such fine-grained information present in lower resolution images by exploiting the micro-feature-selective biases of single voxels that stem from the variability of the distribution of cortical columns or their vascular architecture (Beeck, 2010; Freeman et al., 2011; Kamitani & Tong, 2005; D J Mannion et al., 2009; Damien J Mannion et al., 2015; Sasaki et al., 2006).

Inspired by the demonstrated fine sensitivity of MVPA to finer scale spatial information, here we apply multivariate analysis to BOLD time-courses in order to maximize sensitivity to neuro-temporal information. Capitalizing on the growing interest surrounding the temporal domain of fMRI, we propose a method that captures the temporal characteristics of the BOLD signal at the multi-voxel pattern level. The method, first introduced in Ramon et al. (Ramon et al., 2015), consists of probing single trial events to investigate how the associated representational pattern of activity (Kriegeskorte et al., 2008; Kriegeskorte & Kievit, 2013) for a given stimulus evolves over time. This enables the creation of Single Trial Representational Dissimilarity Matrices (stRDMs), which allows assessing the temporal evolution of the (dis)similarity of these activity patterns.

As previously shown on real data (Ramon et al., 2015), here we demonstrate on synthetically generated data that our approach can detect multivariate differences over time in the *absence of univariate* amplitude modulations across conditions. As such, our temporal multivoxel pattern analysis (tMVPA) offers a different albeit potentially complementary approach to examining BOLD temporal dynamics. We further present a sliding window statistical analysis of these stRDMs that allows quantifying the precise temporal window displaying the effect of interest. We estimate the power and sensitivity of the technique using Monte Carlo simulations.

## Methods

### Procedure and MRI acquisition

Note that the acquired data were used as a starting point to generate synthetic data with realistic signal properties. Thus, within the context of this paper, the original purpose and the hypothesis of the experiment are irrelevant.

#### Participants

20 healthy right-handed subjects (age range: 18-31) participated in the study. Of these, 10 were WC (5 females; mean age, 24) and 10 were EA (4 females; mean age, 22). Three participants (1 WC 2 EA) were excluded from the analysis due to excessive motion during scanning (details below). All subjects had normal, or corrected vision and provided written informed consent. The ethical committee of College of Medical, Veterinary and Life Sciences at the University of Glasgow approved the experiments.

#### Stimuli and procedure

The experimental procedure consisted of a standard block design face localizer and a simple slow event-related face paradigm. All visual stimuli used for the face localizer consisted of front-view gray scale photographs depicting 20 different faces (5 identities × 2 genders × 2 races, taken from the JACFEE database (Matsumoto, D., & Ekman, 1988)), houses (Husk et al., 2007) and textures of noise, respectively. Noise texture stimuli were created by combining the mean amplitude spectrum across faces and houses with random phase spectra sampled from a Gaussian distribution, thereby lending them to contain the same amplitude spectrum as the face and house stimuli. For the main slow event-related experiment, a different set of images used in previous studies (Michel et al., 2006) was utilized which also consisted of 20 front-view gray scale photographs of WC and EA (again 5 identities × 2 genders × 2 races). All images subtended approximately 3.75 × 4.25° of visual angle. Face stimuli were cropped to remove external features; none had particularly distinctive features and male faces were clean-shaven. The stimuli were centered in a 52 × 52 cm background of average luminance (25.4 cd/m2, 23.5-30.1). All images were equated in terms of luminance, contrast and spatial frequency content by taking the average of the amplitude spectra of all stimuli and combining that average spectrum with the original phase spectra to reconstruct each individual stimulus. The root mean square contrast (i.e. the standard deviation of the pixel intensities) was also kept constant across stimuli. Stimuli were projected from the back of the scanner on a round screen situated in the scanner tunnel and occupying the whole width of the tunnel (i.e. 60 cm of diameter). Participants viewed the images through a mirror placed on the head coil.

All participants completed two runs of the block design face localizer fMRI experiment to define the areas responding preferentially to faces (∼12 min/run), and three runs of the main event-related design experiment aimed at measuring the neural activity elicited by individual SR and OR identities (∼16 min/run).

#### Face localizer

Face localizer runs involved presentation of blocks of WC or EA faces, houses and noise textures. Each run began with presentation of black fixation cross displayed on grey background for 20 sec and consisted of 24 randomly presented blocks of images. Each block (6 blocks/category; separated by a 12 sec fixation) involved presentation of 10 different stimuli randomly presented for 800 ms, separated by a 400 ms ISI. To minimize attentional confounds on the BOLD signal related to the race of the stimuli, we implemented an orthogonal task. Participants were instructed to respond to red or green stimuli which (10% of the images, i.e. one red or green stimulus per block), by pressing a button on a response pad held in their right hand.

#### Event-related experiment

Each run of the event-related face experiment began and ended with 20 seconds fixation and consisted of 80 events (10 identities per race x 2 races x 4 repetitions per identity). Face stimuli were displayed for 850 ms followed by a 11.15 sec fixation cross; participants were instructed to maintain fixation on a central fixation cross throughout each 12 sec event. As for the face localizer scans, an orthogonal task was employed with participants responding to a change in the color of the fixation cross (red or green, for 200-1200 ms at a random time within an event, before reverting to its original color) by pressing a button.

#### MRI acquisition protocol

All MRI data were collected with a 3-T Siemens Tim Trio System with a 32-channel head coil and integrated parallel imaging techniques (IPAT factor: 2). Functional MRI volumes were collected using an echo-planar acquisition sequence [*localizer runs*: repetition time (TR), 2000 ms; echo time (TE), 30 ms; field of view (FOV), 210 × 210 mm; flip angle (FA), 77°; 36 axial slices; spatial resolution, 3mm isotropic voxels; *event-related runs*: TR, 1000 ms; TE, 30 ms; FOV, 210 × 210 mm; FA, 62°; 16–18 axial slices; spatial resolution, 3 × 3 × 4 mm voxels]. Slices were positioned to maximize coverage of occipito-temporal regions. T1-weighted anatomical images were obtained using an MPRAGE sequence (192 slices; TR, 1900 ms; FOV, 256 × 256 mm; flip angle, 9°; TE, 2.52 ms; spatial resolution, 1 mm isotropic voxels). For participants who were re-scanned due to movement artifacts, separate anatomical scans were recorded for each scanning session to facilitate realignment of the functional data.

#### MRI data preprocessing

fMRI data were preprocessed in native space using BrainVoyager QX version 2.1 (Brain Innovation). Functional images were slice-scan time corrected, three-dimensional motion corrected with reference to the functional volume taken just before the anatomical scan, high-pass filtered using a Fourier basis set of three cycles per run (including linear trend). Images were co-registered with the anatomical set and spatially normalized into Talairach space (Talairach, J., & Tournoux, 1988); images from localizer runs were spatially smoothed with a full-width at half-maximum of 4 mm.

#### Functional ROI definition

Five functional ROIs were identified from the localizer runs. Individual participants bilateral FFA, bilateral OFA, and right AIT were identified by performing F-tests on all the voxels in the brain and determining the peak voxel of the activation clusters identified by the contrast (WC + AC) faces > (Houses +Noise) located in the bilateral fusiform and inferior occipital gyrus, respectively. To control for type I errors, False positive Discovery Rate (FDR) was implemented as a multiple comparison correction. The significance threshold was set to *q<.05* for all ROIs and participants. The corresponding masks for these ROIs were exported into MATLAB (MathWorks) for subsequent analyses. Across all participants from both groups (WC and EA), we identified 86 ROIs in total. While bilateral FFA and right OFA were identified in all participants, a few subjects did not have a clear definition of left OFA and right AIT. The average number of voxel across all ROIs was 47.9 (std: 16.7).

#### BOLD percent signal change and epochs definition

For each voxel, we computed BOLD percent signal change by dividing the raw BOLD time course by its mean. We then defined the epochs of interest as those portions of the whole BOLD time series ranging from 1 TR prior to 14 TRs after stimulus onset. For each single trial we extracted these 15-TR long time-courses from all the voxels within each ROI of every subject. These BOLD percent signal change epochs were saved as a matrix that we used to generate synthetic data using Monte Carlo simulations (details below).

### Temporal multivariate pattern analysis (tMVPA)

In this paper, we developed a novel multivariate temporal analysis for the BOLD time-course, inspired by representational similarity analysis (Kriegeskorte & Kievit, 2013). This approach assesses the temporal evolution of the degree of dissimilarity of neural representations - defined as the pattern of BOLD response across all voxels - elicited by different time points (Ramon et al., 2015). It involves computing Single Trial Representational Dissimilarity matrices (stRDMs) within a selected ROI between two conditions (e.g., *baseline* and *treatment* condition). We compute stRDMs on the BOLD percent signal change independently per subject and condition as follows: for each condition, we iteratively correlated (Pearson r) the values of all the voxels at one time point with all the remaining ones amongst the epochs of two different trials (e.g. the time course elicited by trial 1 and that elicited by trial and calculated the correlation distance (i.e. 1-r; see Figure 1). This procedure was repeated across all possible trial pair combinations. The resulting matrices were fisher-z transformed to render the skewed Pearson-r distribution approximately normal. We then averaged (10% trimmed mean) the single trial correlational distance matrices to obtain the single subject stRDM.

**Figure 1.**
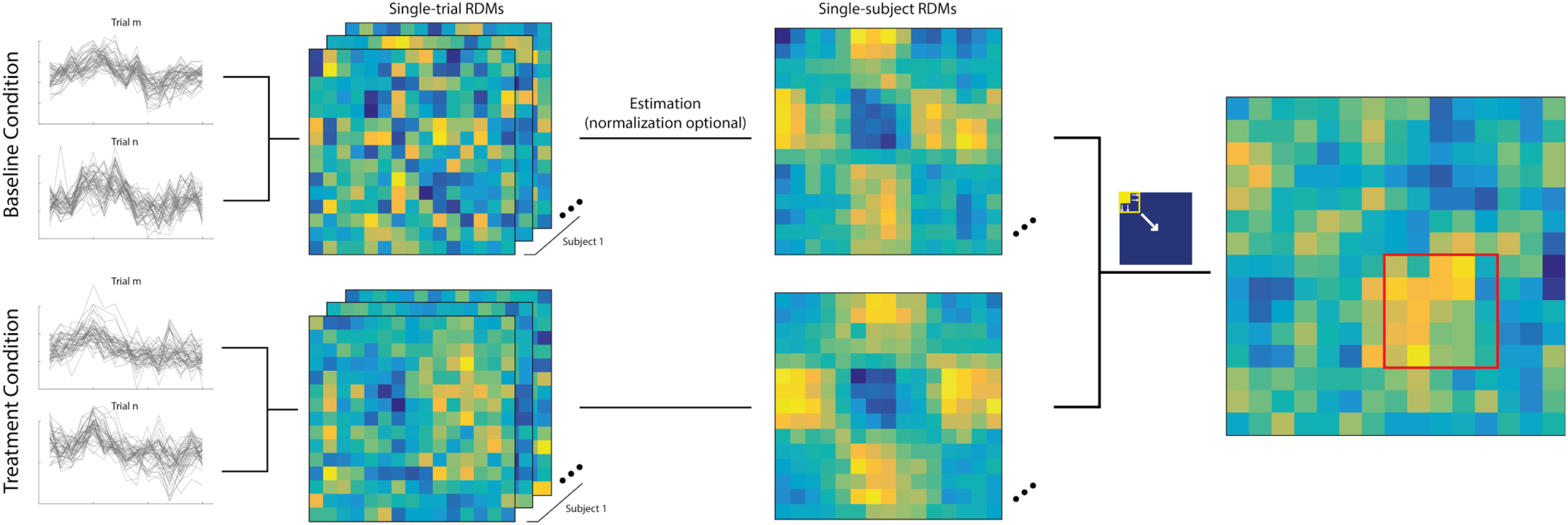
The temporal multivariate pattern analysis (tMVPA) procedure. Cool colors indicate higher similarity between neural representations elicited by any 2 given time points. Warm colors indicate higher dissimilarity or distinctiveness amongst neural representations. Each row and column represents a single TR.

While all subsequent statistical analyses were performed on the fisher-z transformed values, for visualization purposes (figure 1) and to render the values within the stRDMs interpretable, we performed the inverse of the fisher-z normalization on the final averaged stRDM.

To test for statistically significant differences between the stRDMs from different conditions (i.e., *baseline* and *treatment* condition), we implemented an expanding sliding window approach. We started by computing a simple subtraction between the stRDMs of the 2 conditions of interest. We then centered a 2×2 pixel window (figure 2) on the first point of the diagonal of the matrix. We then computed the 10% trimmed mean across the values within the window. We divided this mean by the standard error of the values within the window. Given that the standard error is a function of the variance weighted by the number of data points, this procedure was implemented to partially account for the relative difference in terms of data points and variance across windows of different sizes. We then performed (1-*alpha*) bootstrap confidence interval (CIs) analyses by sampling subjects with replacement 500 times. Importantly, we adjusted the threshold (*alpha* above) for determining high and low CIs as a function of the total number of windows to account for multiple comparison problems (i.e. Bonferroni correction). The analysis was repeated on increasingly larger windows that expanded by 1 pixel in each direction (when applicable), centered on each point of the diagonal (figure 2). Differences between conditions were inferred when the btCIs did not include zero. This expanding sliding window approach allows investigating whether potential differences across stRDMs encompass a few time points or whether these are sustained over a larger time window.

**Figure 2:**
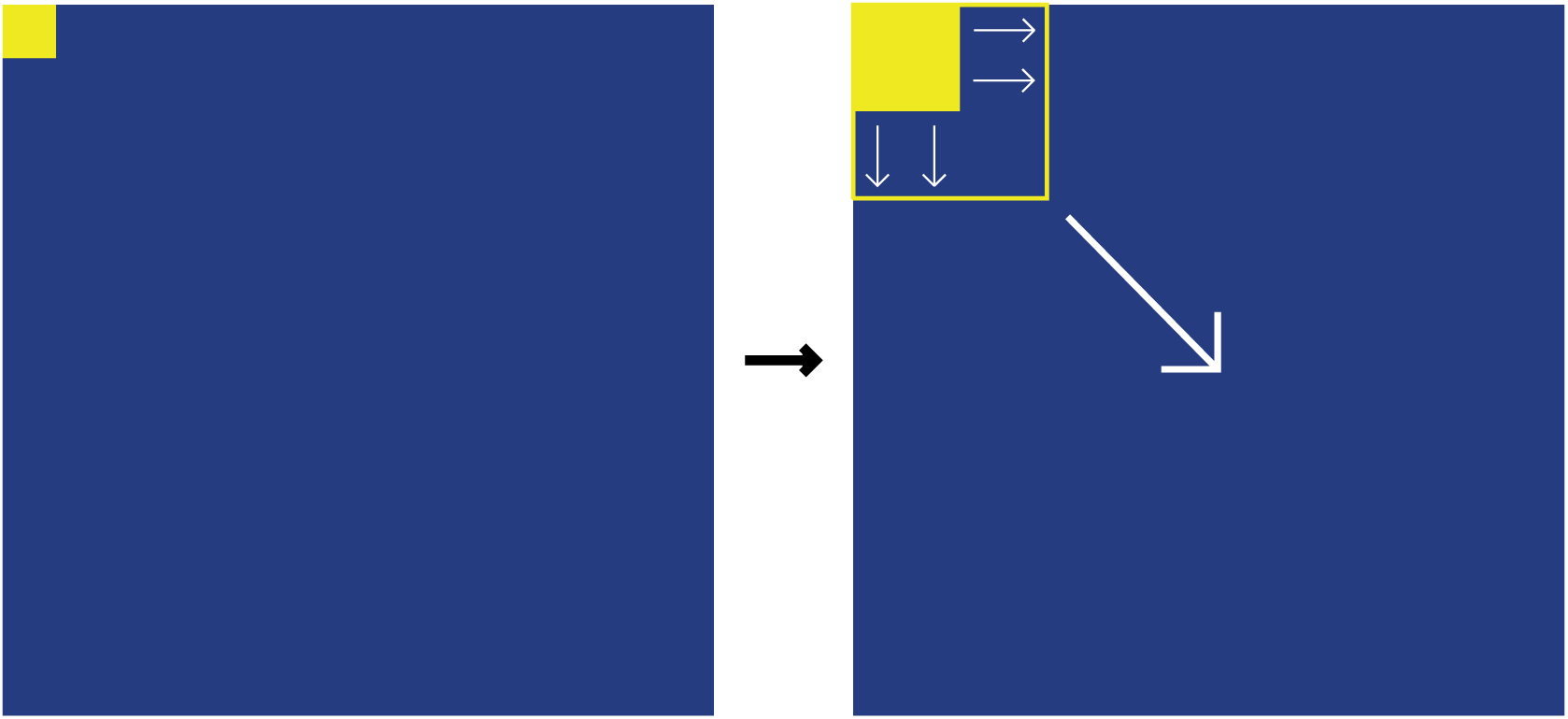
Expanding Sliding Window approach. The panel on the right depicts the starting window size and location, while the panel on the right represents this same window “expanding” (as indicated by the thin pairs of white arrows) and sliding (as indicated by the larger arrow).

### Synthetic Data Generation and Validation

The following sections describe the procedure we implemented for the synthetic data generation process and the approach we adopted to estimate the power and Family-wise error rate (FWER) of our proposed multivariate temporal analysis. In brief, we employed Monte Carlo (MC) simulation on synthetic data to estimate the FWER and the statistical power of our proposed method, explicitly manipulating a number of parameters (see the *Manipulated parameters paragraph*). In order to reproduce realistic fMRI noise and signal properties, we generated synthetic data starting from the BOLD signal recorded during the event-related experiment. We created a complete dataset comprised of 2 conditions (i.e. *Baseline* and *Treatment*). Importantly, we generated Baseline and Treatment conditions under 2 distinct scenarios: 1) under H0 (i.e. no multivariate differences between conditions), thus being in the ideal context to measure our approach’s FWER, as any statistical difference detected by our approach would be a false positive; and 2) under H1 (i.e. artificially introducing multivariate pattern differences between conditions - see s*ynthetic multivariate effect*) to test our approach’s power (see below for more details).

#### Synthetic data generation

Starting from the single trial BOLD time course matrix (see the *BOLD percent signal change and epochs definition* paragraph), we extracted single trial epochs from one of the 20 conditions for one participant across one run and using just a single ROI. We saved the extracted BOLD values in a 3D *Raw_singletrials_BOLD* matrix with dimensions <u>[number of trials * number of voxels * number of time points]</u>. From the *Raw_singletrials_BOLD* matrix we calculated the mean and the variance across voxels, and then saved these 2 metrics in 1D vectors of size <u>[number of time points].</u> We refer to these vectors, representing respectively the average HRF for a given ROI and the voxel-wise variance within that same ROI, as *mu_BOLD(time point)* and *var_BOLD(time point)*. We then calculated the residual between the single trials epochs and their mean (across trials) for each voxel and time point, and then saved these values in a [<u>number of trials * number of voxels * number of time points], a</u> 3D matrix that we refer to as *sigma_BOLD(trial, voxel, time point)*).

We repeated the procedure described above for all conditions, runs, ROIs, and subjects. The resulting *mu_BOLD, var_BOLD*, and *sigma_BOLD* were flattened and saved in 2 dimensional matrices: **E, V,** and **S**. Note that the matrices **E, V,** and **S** have an equal numbers of columns, corresponding to the number of time points per epoch of interest (i.e. 15), but a different number of rows. For the matrices **E** and **V**, containing, respectively, the mean time courses across voxels and the variance across voxels, the number of rows was equal to [<u>number of subjects * number of runs * number of conditions * number of ROIs]</u>; while the number of rows for matrix **S**, containing the single-trial residual for each voxel, was equal to [<u>number of subjects * number of runs * number of conditions * number of ROIs * number of trials * number of voxels per ROI]</u>.

The raw BOLD signal was thus fully represented in matrices **E, V,** and **S**. To generate synthetic data for one subject we randomly sampled one row vector from **E** and **V** and generated a 2D <u>[number of voxel * number of time points]</u> matrix, representing the mean (across trials) time course for all voxels within a given ROI. We then injected the trials’ variation from their mean by randomly sampling from **S (see below for details)**.

In order to generate the Baseline and Treatment conditions, we implemented very similar, albeit slightly different procedures. The first step of the data generation process (step 0) was the same regardless of the generation goal. For each MC simulation, we began by randomly selecting a row vector ***e*** from matrix **E**, representing the group average time course for a hypothetical ROI.

For the Baseline condition, independently per subject we generated a <u>number of voxels</u> (*nv*) <u>number of time points</u> (*ntp*) <u>* number of trials</u> (*ntrial*) matrix **MB**, following the 9 step algorithm below:

- Step 1, we randomly selected a row vector ***v*** from matrix **V**, and *nv*ntrial* rows vectors from matrix **S** to get ***sv***. To have full control of the simulation study, we kept the variance across time points within a single voxel and a single trial constant by setting *v_1* = *v_2* = … = *v_ntp* = *mean(**v**)* and *sv_1,i* = *sv_2,i* = … = *sv_ntp,i* = *mean(**sv**) for i ∼ [1, nv]*.
- Step 2, we repeated *nv* copies of array ***e*** and transformed them into a *nv*ntp* matrix **ev**.
- Step 3, we repeated *nv* copies of array ***v*** and transformed them into a *nv*ntp* matrix **vv**.
- Step 4, we generated a *nv*ntp* matrix **dv1** to represent the variance across voxel. Each element in **dv1** was generated following one of 3 distributions: either *Normal(mu=0, sd=1), Uniform 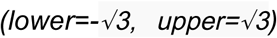,* or *Exponential(lambda=1) - 1*. These three distributions all have mean equal to 0 and variance equal to 1.
- Step 5, the mean BOLD time course for each voxel **Mp** was generated following the equation:

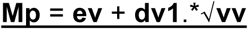 Where “**.\***” indicates the element-wise multiplication. By doing this, **Mp** satisfies *mean(***Mp***) =* ***e*** and *var(***Mp***) =* ***v***. **Mp** is an *nv*ntp* matrix representing the single voxel BOLD time course.
- Step 6, we repeated *ntrial* copies of matrix **Mp** and transformed them into a *nv*ntp*ntrial* matrix **MP**.
- Step 7, we reshaped the residual matrix ***sv*** into an *nv*ntp*ntrial* matrix and computed the variance across trials. The resulting *nv*ntp* matrix was then repeated and reshaped into an *nv*ntp*ntrial* matrix ***svt*** representing the single trials residuals for each voxel and timepoint.
- Step 8, we generated an *nv * ntp*ntrial* matrix **dv2**. Similar to **dv1**, each element in **dv2 h**followed one of 3 distributions: either *Normal(mu=0, sd=1), Uniform 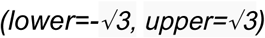,* or *Exponential(lambda=1) - 1*. **dv2** represents the noise at the single trial level for each voxel.
- Step 9, finally, we computed the single trials BOLD time course matrix **MB** following the equation:

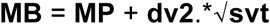 Notice that the mean and variance across trials for **MB** satisfies *mean(***MB***) =* **Mp** and *var(***MB***) =* ***svt***.

These 9 steps were repeated for all subjects.

Similar to the baseline conditions, we generated an *nv * ntp*ntrial* **MT** Treatment condition matrix for each subject following the same 9 steps.

When no effect was introduced in the Treatment condition (i.e. FWER estimation, see below), the **MT** matrix creation began directly at step 7 (through to 9), starting from the same **MP** and ***sv*** generated for the Baseline condition using steps 1 to 6. Thus, the **MT** mean and variance across trials satisfies *mean(***MT***) =* **Mp** and *var(***MT***) =* ***svt***.

#### Synthetic multivariate effect

Our procedure to introduce multivariate differences between the baseline and treatment conditions consisted of rendering the voxel response for some selected time points in the treatment condition highly correlated across trials. To achieve this, we first repeated steps 1 to 9 to generate matrix **Mp’**, containing the treatment condition mean BOLD time course across all trials for all voxels within a given ROI; **MP’,** containing the single trials BOLD time-course for all voxels within any given ROI; **svt’,** containing the residuals between the single trials and average across trials for each voxel, time-point, and trial; and **MT’**, containing the single trials’ BOLD time courses for all voxels within a given ROI. We therefore modulated *k* consecutive time points in matrix **MT’** to introduce correlation in the synthetic signal by rotating the data matrix **MT** to reduce the multivariate distance across trials^1^. Independently for each of the *k* time points, we first repeated step 8 to generate a new independent and identically distributed (i.i.d.) noise matrix **dv2’**. We then computed the BOLD time course for the treatment condition **MT** following the equation:

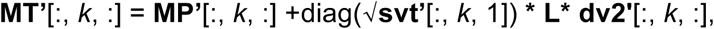

where **L** is the Cholesky factor of a correlation matrix randomly sampled from a LKJ correlation distribution (Lewandowski et al., 2009). Therefore, the variance across voxels for *k-th* time points of some selected voxels was identical:

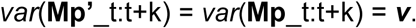

Notice that the univariate pattern in **Mp’** was kept constant: *mean(***Mp’***) =* ***e***. Moreover, the mean and variance across trials for **MT’** also satisfies *mean(***MT’***) =* **Mp’** and *var(***MT’***) =* ***svt’***.

The resulting data matrix **MB** and **MT’** represented the full synthetic dataset for one subject. We repeated the above 9 steps to generate *k* (number of subject) **MB** and **MT’** matrices. We therefore implemented our TMPVA analysis to test for multivariate differences between the treatment and baseline conditions. We repeated this MC simulation 1000 times for each combination of parameters (details below).

#### Manipulated parameters

In an attempt to maximally parameterize our validation procedure while keeping within the boundaries of reasonable computational demands, we manipulated the following 4 parameters: 1) number of trials per condition, 2) number of subjects per group, 3) number of time points at which the effect was introduced, and 4) the percentage of subjects (or trials for the single subject validation procedure) in which the effect was introduced (i.e. the target power).

1. the number of trials varied across 4 different levels: 4, 8, 12, and 16.
2. for the number of subjects, we tested 4 sample sizes: 6, 10, 14, 18 participants.
3. while the multivariate effect always began at TR 5, the number of time points at which the effect was introduced varied across 4 different levels: 2, 3, 4, 5.
4. the percentage of sample showing effect (i.e. power), varied across three different levels: 50%, 65%, and 80%.

Additionally, the number of voxels (range [30, 60]) per simulated subject was randomized across all MC simulations. We thus ran independent MC simulations for all possible combinations of the different parameter levels. This parameterization of the MC simulation was implemented to evaluate the reliability and sensitivity of our method in different experimental contexts. Note that we introduced a multivariate effect for our power analysis at time point 7 (up to time point 11, depending on the number of manipulated time points). For the estimation of FWER, only number of trials and number of subjects were relevant parameters. For each unique parameter combination, we computed 95% bootstrap CI based on 500 bootstraps, and repeated this procedure 1000 times.

Importantly, we validated our tMVPA approach within two different settings: *group analysis* and *single subject analysis*. In the group analysis setting, to manipulate the target power we varied the percentage of subjects in which we introduced correlation across voxels (i.e. the synthetic multivariate effect). In the single subject validation setting, the target power was instead manipulated by varying the percentage of trials in which the multivariate pattern was introduced (i.e. 50%, 65% or 80% of the trials).

#### FWER estimation

To estimate the FWER, we performed tMVPA analysis to test for multivariate differences between the time courses of the baseline and treatment conditions, prior to introducing correlation across voxels at selected time points. We thus counted the number of significant events detected by our approach. We repeated this procedure 1000 times. Since baseline and treatment conditions were created under H0 (i.e. no differences between them), significant differences detected by our approach were considered to be false positives (i.e. type II error). The FWER was thus computed as the total number of significant time windows divided by 1000 (i.e. the total number of MC simulation).

#### Statistical power estimation

For statistical power estimation we, instead, generated 1000 *treatment* conditions following a procedure similar to the generation of the *baseline* condition (i.e. steps 1 to 9 as described earlier). We additionally introduced multivariate differences between conditions (see *Synthetic multivariate effect*) in a number of subjects by manipulating the pattern of voxels within a given ROI over some selected time points (see *Manipulated parameters for more details*). Importantly, no univariate differences (i.e. no differences between the time courses averaged across voxels - see figure 3 and 4) between the two conditions existed over these time points. The target power of the tMVPA approach was represented by the percentage of subjects for whom we introduced multivariate differences between conditions. For example, if we introduced correlation across voxels in 80% of the subjects, we expected the tMVPA to report significant differences 80% of the time across all simulations where the effect was introduced. The statistical power of tMVPA was thus computed as the total number of significant time windows detected divided by the total number of MC simulations.

**Figure 3:**
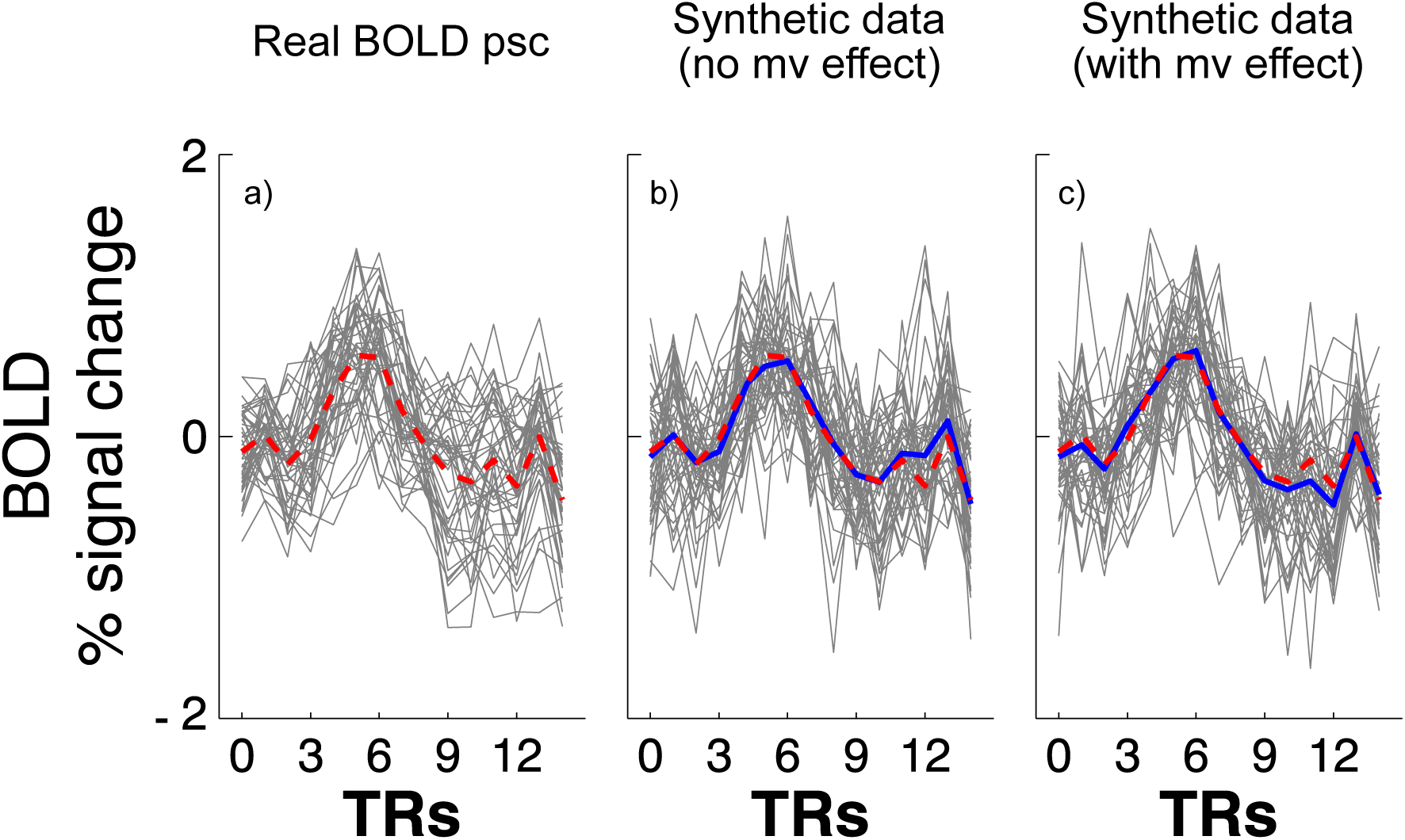
Panel a) portrays an example of real BOLD percentage signal change (psc) time course for all voxels in a given ROI for a single subject. The grey line plots show the BOLD time course for each voxel, while red dashed line shows the average BOLD time course. Panels b) and c) depict the generated synthetic BOLD time course created using the same mean and variance of the real BOLD time course. Panel b) shows an example of the synthetic baseline condition - i.e. no multivariate (mv) effect; and Panel c) shows an example of a synthetic treatment condition where we introduced a mv effect (see the Synthetic multivariate effect paragraph) over time points 5-7. Grey line plots show single voxels, the red dashed line shows the average time course of the real signal, the blue line shows the average time course of the synthetic data.

**Figure 4:**
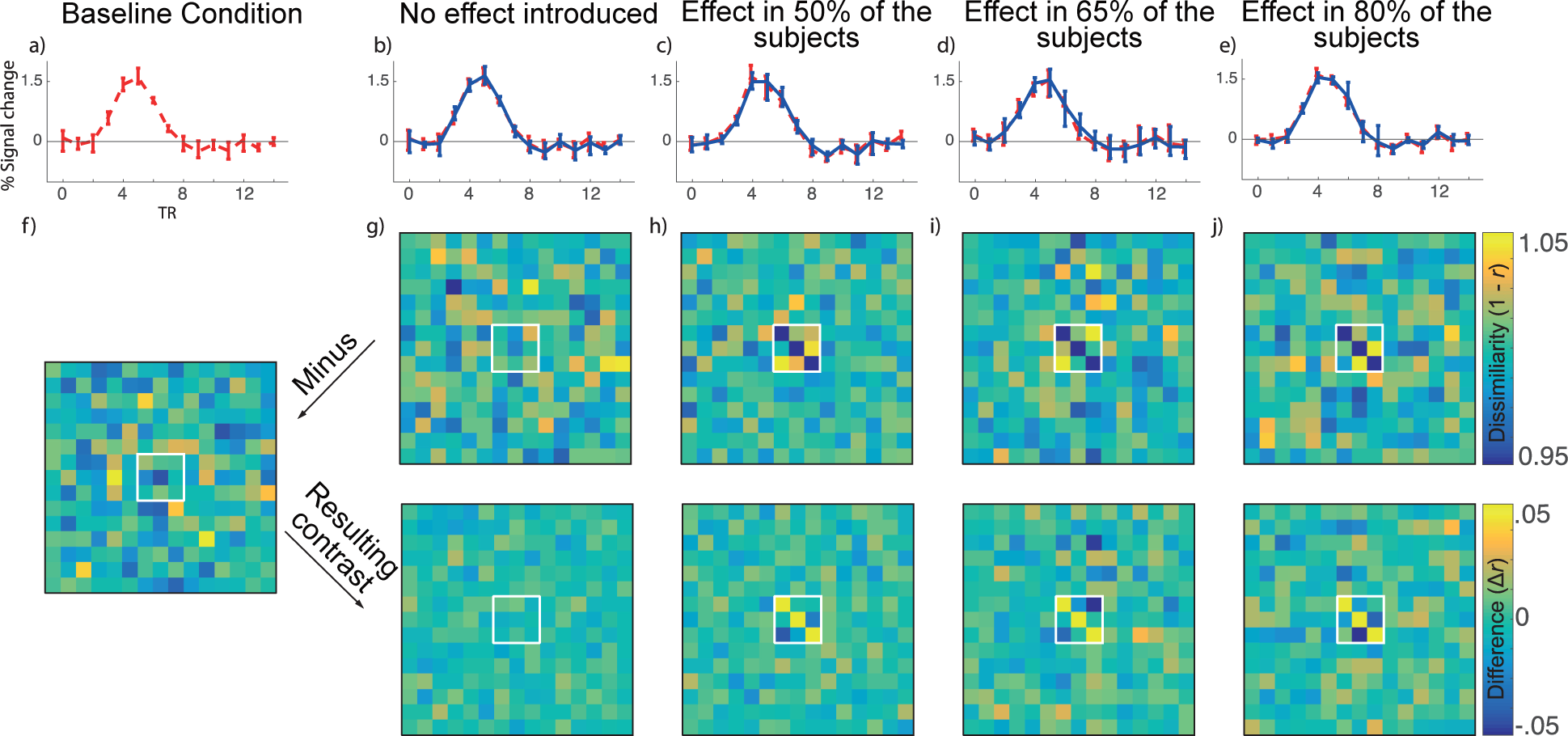
Synthetic data for the 18 subjects and 14 trials group a)-e) Average time course across voxel participant within a ROI. Red line shows baseline condition (a) and blue line shows Treatment condition. Error-bars shows 95% bootstrapped confidence interval across subjects for each time point. f) stRDM of the baseline condition. g) stRDM of the treatment condition when no effect is introduced (to estimate FWER). h)-j) stRDM of the treatment condition when different strengths of the multivariate effect is introduced over time-points 5-7.

## Results

95% bootstrap confidence intervals (btCIs) computed across our MC simulations showed that manipulating the number of time points at which we introduced the synthetic multivariate effect did not significantly (p>.05) impact FWER and power estimations (see supplementary section). Additionally, we observed that the distribution from which we sampled the synthetic noise did not significantly (p>.05) modulate FWER and power estimations (see supplementary section). We therefore only report the results for synthetic data with a multivariate effect over 3 time-points, generated by sampling noise from a normal distribution. Figures and results for the remaining levels of these 2 parameters as well as detailed tables reporting mean and bootstrap CIs can be found in the supplementary section.

In the following paragraph we report the mean across all MC simulations and standard deviation (std) of the peak amplitude of the BOLD % signal change time course. We further report the mean std across voxels, trials, and time course. In the MC simulations for the group study, the mean peak amplitude (across subjects and MCs) of the generated synthetic BOLD % signal change was 1.222 (std = .531), while a mean std across time 0.353 (std = .137). Moreover, the average std across voxels was 2.815 (std = 2.643) and the average std across trials 1.343 (std = .348). As for the MC simulation for the single subject study, the generated synthetic data set had a mean (across MCs) peak amplitude of 1.247 (std = .533), with a mean std across time 0.357 (std = .106). The mean std across voxels was 2.996 (std = 2.409), and the mean std across trials was 1.364 (std = .362).

Figure 4: (panels a through e) shows the BOLD time course of our synthetic data for the 18 subjects and 16 trials scenario. Error bars represent the 95% bootstrap confidence intervals (btCIs). We infer robust statistical significance (p<.05) when the error bars do not overlap. Our analyses revealed no significant univariate amplitude differences across the whole time course between the *baseline* (red line) and the *treatment* (blue line) conditions for all the parameter manipulations (see manipulated parameters). Importantly, this absence of univariate amplitude differences persisted even after we synthetically introduced multivariate effects at selected time-points. Our tMVPA approach, thus, crucially revealed robust *genuine* multivariate differences across conditions that are not evident in univariate amplitude differences. Note that the introduced multivariate effect is visible by computing the stRDM, as shown in Figure 4f for the *baseline* condition and Figure 4h-4j for the *effect* condition.

### Family-wised error rate (FWER) under H0

For both the group and single subject scenarios, to estimate the FWER we computed the frequency of significant outputs detected by our approach across MC settings, before introducing the multivariate effect. As explained earlier, prior to introducing correlation across voxels over a number of selected time points, we generated the synthetic baseline and treatment data under H0 (i.e. no differences between conditions). We were, therefore, in the ideal context to estimate FWER, as statistically significant differences between conditions were mere type I errors.

#### Group-level analysis

95% bootstrap confidence intervals (btCIs) show that FWERs were significantly below.05 in all MC simulation with sample size > 6 (Figure 5). For N=6 the mean of the estimated FWER was, instead, consistently above .05 (mean FWER: .058), regardless of the number of trials. The 95% btCIs (mean btCIs [.044 .073], however, indicated that even for N=6, FWER are not significantly larger than .05 (see figure 5). While according to Westfall and Young (1993) this still suggests the group analysis is valid, we would recommend caution using our tMVPA with only 6 subjects. This is because the FWER for N=6 were significantly larger than those estimated for all other sample size (6 subjects simulation lowest mean FWER and btCIs: .056; [.042 .07]; highest FWER and btCIs across the remaining MC simulations: .03; [.02 .042]. For a complete table of all FWER and btCIs see supplementary section). **Overall, our approach achieved the desired FWER at 5% under the group analysis setting.**

**Figure 5:**
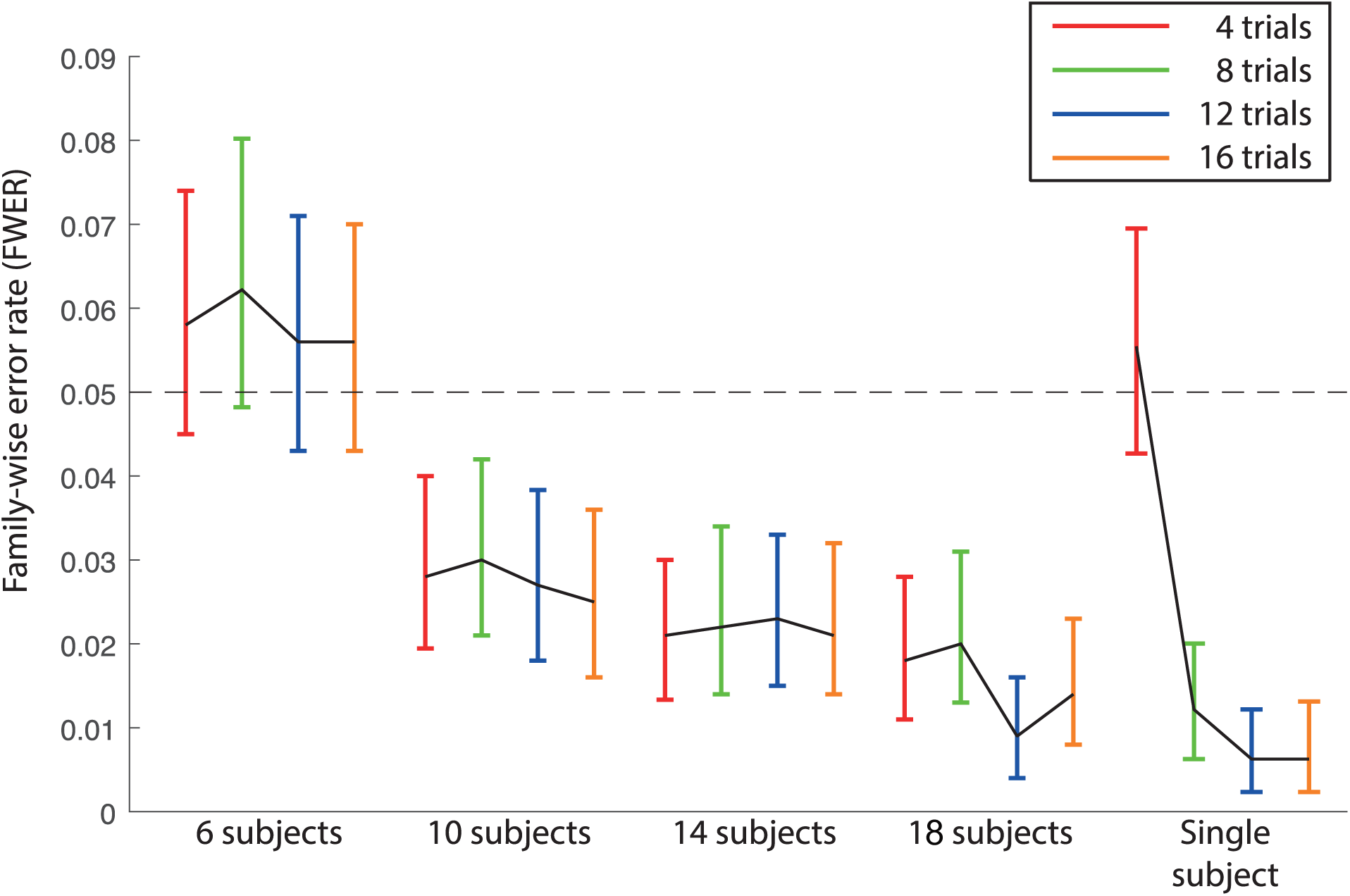
FWER for all trials numbers, subjects groups and for the single subject scenario. Here we show the family-wise error rate for the Monte Carlo simulated synthetic data with noise sampled from a Normal distribution. Error-bars represent the 95% bootstrap confidence interval of the Monte-Carlo simulation.

#### Single-subject analysis

Similarly, FWERs were not significant above .05 in all MC simulations, regardless of the number of trials, as shown in Figure 5 above. The highest FWER is 0.056 [0.043, 0.071] in the simulation with 4 trials, and the lowest FWER is 0.006 [0.003, 0.013] in the simulation with 16 trials (For a complete table of all FWER and btCIs see supplementary section). The 4 trials scenario produced significantly higher FWER than all other trials groups. While still not significantly larger than .05, we would still recommend caution if implementing our TVMPA approach with less than 8 trials, due to the risk of incurring Type I errors. Overall, **the simulation result clearly showed that in a single-subject analysis setting, our approach achieved the desired FWER at 5% even with as little as 8 trials.**

### Power analysis

#### Group-level analysis

btCIs analysis generally revealed that for the group scenario, regardless of the number of trials, tMVPA was relatively underpowered when differences across conditions were present in 50% and (to a lesser extent) 65% of the subjects. As shown in Figure 6, the power of our approach increases as the number of subjects and the number of trials increases. With the effect introduced in 50% of the subjects, we estimated a power of 0.15 [0.132, 0.176] at the lowest number of subjects and trials (6 subjects with 4 trials each), to 0.27 [0.247, 0.297] at the highest tested number of subjects and trials (18 subjects with 16 trials each). Importantly, when we introduced the effect in 80 % of the subjects, the 16 trials simulations led to significantly (p<.05) higher power than the 8 and 4 trials scenarios for all sample sizes. Moreover, while generally displaying higher mean power, the 16 trials simulation never significantly (p>.05) differed from the 12 trials one. It is also worth noting that when N = 18, both the 16 and 12 trials simulations led to significantly higher power (p<.05) compared to the 4 and 8 trials simulations, regardless of the number of subjects in which we introduced an effect. Furthermore, for the 14 subjects simulations only, the power estimated for the 4 trials scenario was significantly lower than all other group sizes, regardless of the number of subjects displaying the effect.

Not surprisingly, the highest statistical power was reached in the 18 subjects simulations with a minimum of 12 trials. Within this context, the tMVPA approaches 0.8 when we introduced the multivariate effect in 80% of the subjects (0.76 [0.731, 0.784], see also Figure 6). A detailed report of mean power and btCIs for all MC simulations can be found in the supplementary section.

**Figure 6:**
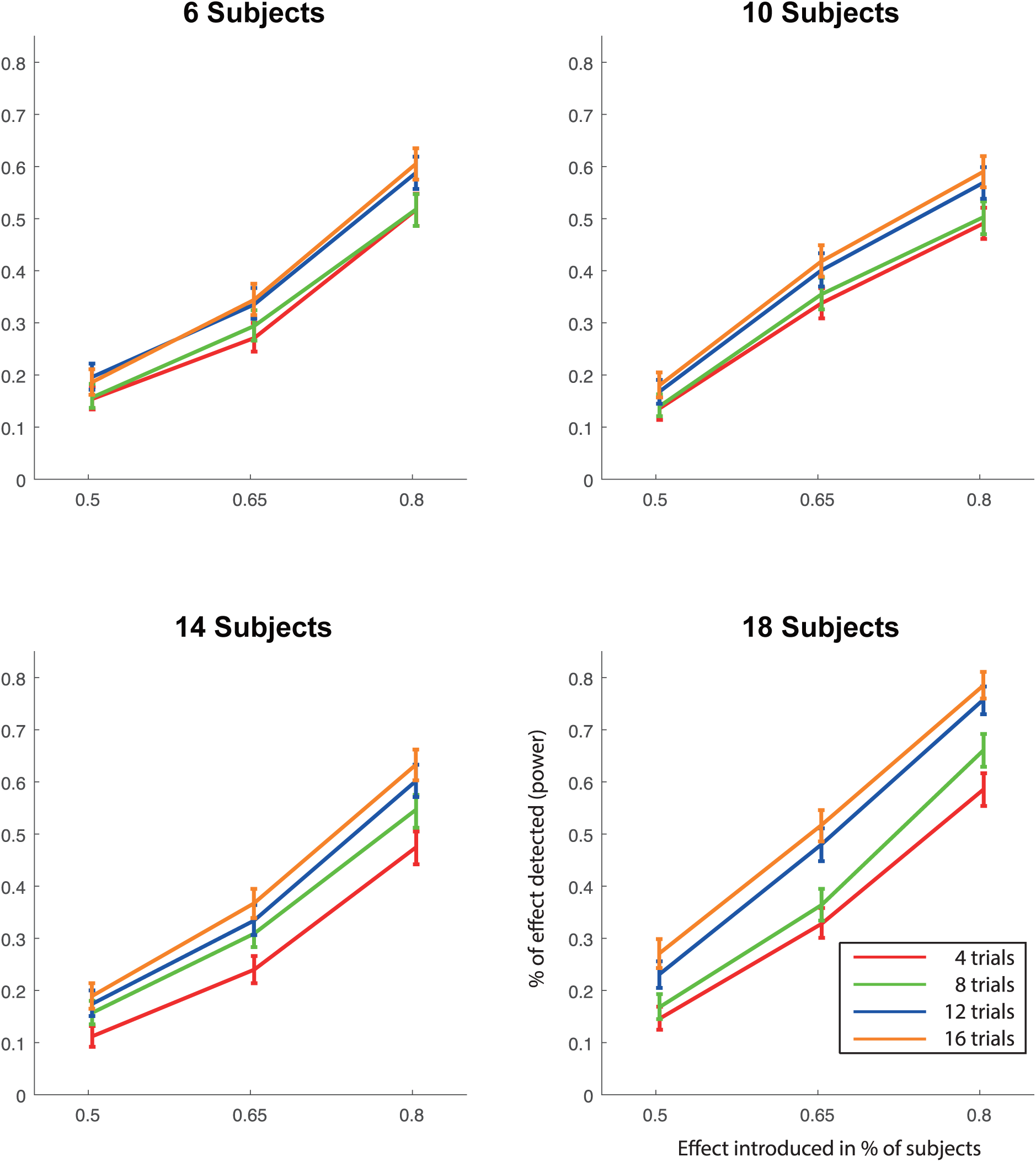
Statistical power of the group-level analysis. Error-bars represent the 95% bootstrapped confidence intervals across Monte-Carlo simulations.

#### Single-subject analysis

As shown in Figure 7, the power of our approach increases as the number of trials increase. With the effect introduced in 50% of the trials, we estimated the power of our proposed approach at 0.10 [0.087, 0.126] with 4 trials, and at 0.28 [0.260, 0.317] with 16 trials. With the total number of 16 trials, the statistical power of the proposed approach reached 0.8 when the effect was introduced in 80% of the trials (0.80 [0.776, 0.827]). Importantly, regardless of the percentage of trials in which we introduced the effect, the 16 trials simulations led to significantly (p<.05) higher power compared to all other simulations. Moreover, while significantly (p<.05) lower than its 16 trials counterpart, the 12 trials simulations also led to significantly (p<.05) higher power than the 2 remaining trials scenarios (see figure 7), peaking when 80% of the trials showed a multivariate effect (mean power: 0.719; btCIs: [0.691, 0.747]). A detailed report of mean power and btCIs for all MC simulations can be found in the supplementary section.

**Figure 7:**
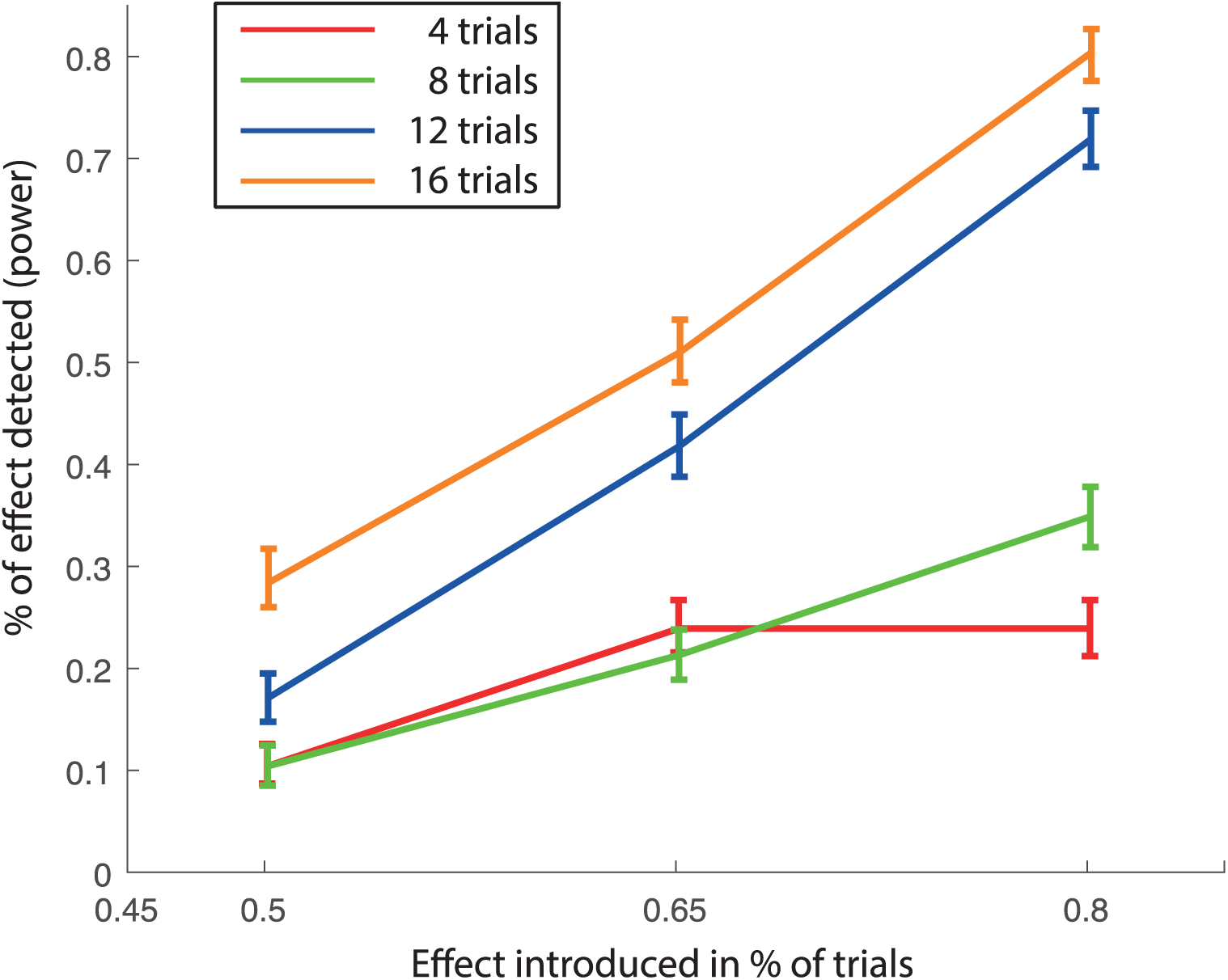
Statistical power of the single-subject analysis. Error bars represent the 95% bootstrapped confidence intervals across Monte-Carlo simulations

## Discussion

In this paper, we present temporal Multivariate Pattern analysis (tMVPA), a method that we developed to quantify the temporal evolution of single trial dissimilarity across multivoxel patterns evoked by a given stimulus within a defined ROI. tMVPA builds upon the generation of single trial Representation Dissimilarity Matrices (stRDM) independently per ROI and condition: for all trials pairs, we iteratively cross-correlate the multivoxel pattern of BOLD % change across all possible time points combinations and we calculate its correlation distance (1-r). We then implemented a robust expanding sliding window approach to identify the temporal loci where statistically significant differences between conditions can be inferred (see methods). We validated this method for group and single subject analyses on data that were synthetically generated using noise (e.g. std across voxels, trials and time points) and signal (e.g. the BOLD time course) parameters derived from real fMRI data. Our validation analysis revealed 2 main findings: 1) our tMVPA approach reached the desired FWER (<=.05) for both the group and single subject approach; and 2) Our power analysis showed that: a) for the *group* scenario, the tMVPA approach reached the desired power with a sample size of 18 subjects, each with 12 trials or more, when 80% of the participants displayed the desired multivariate effect. In all other contexts (i.e. < 18 subjects, < 12 trials and < 80% of subjects showing the effect), our method tends to be relatively underpowered and b) similarly, for the *single subject* scenario, our approach reached the desired power with at least 12 trials, when the multivariate effect of interest was present in 80% of them. All other simulation scenarios failed to reach the target power. These findings are discussed in detail below.

### Group analysis

Simulation results indicate that when the sample size is less than 8 subjects, regardless of the number of trials per condition or percentage of effect introduced, our technique is significantly (p<.05) below the lower margin of the desired FWER (.05) (Figure 7). Thus, a minimum of 8 subjects is needed to maintain Type I error rate. Moreover, it is worth noting that for N=6, FWER is not significantly larger than .05, a finding which advocates the validity of the group analysis (Westfall, P. H., & Young, 1993; Westfall et al., 1993) (at least in terms of false positive rate). Nonetheless, we observe that when N=6, the estimated FWERs are significantly larger than all other samples and MC simulations (see figure 5), which may significantly inflate the occurrence of Type I errors for this specific sample size.

Furthermore, the results of our power analysis suggest that a minimum of 18 subjects with at least 12 trials per condition is required to achieve adequate statistical power. While a sample size of 18 subjects could be regarded as sufficient for the majority of current fMRI studies, low N is considered one of the main culprits for the so called “replication crisis” (Button et al., 2013; Maxwell et al., 2015; Schooler, 2014). Consequently, the field of science as a whole, and specifically disciplines such as psychology and cognitive neuroscience, is undergoing a targeted endeavor aimed at augmenting the experimental sample size, in an effort to increase statistical power and produce replicable results (Button et al., 2013; Maxwell et al., 2015; Schooler, 2014). Within this context, a sample size of 18 participants does not therefore seem prohibitive. Taken together, power analysis and FWER estimation indicate that a minimum of 18 subjects and 12 trials are required to implement tMVPA at the group level.

#### Single subject analysis

One of the main advantages of the tMVPA analysis is the exploitation of single trials in computing temporal RDMs. Generating RDMs by correlating all possible single trial pairs leads to a distribution of single trial RDMs (stRDMs), which allows one to carry out second order inferential statistics at the single subject level. This procedure permits full exploitation of the trial-by-trial variability, which is lost in the group-level approach due to averaging. It is worth noting that, while still not significantly larger than the desired FWER of .05, the single subject validation procedure indicates that the 4 trial scenario produces significantly more FWER than all other trials groups. Not surprisingly, the peak statistical power is achieved for the 16 trials simulations (figure 7). The 12 trials simulations, however, led to significantly higher statistical power than its 4 and 8 trials counterparts. Crucially, when the multivariate effect of interest is present in at least 80% of the trials, our approach achieves the desired power with 12 trials or more. With a minimum of 12 trials across runs, our approach reaches the desired power and FWER. This finding makes our tMVPA appealing and powerful, not only to carry out single subject statistics, but to investigate issues that have thus far been elusive to the world of cognitive neuroscience, such as individual differences in the BOLD response. Moreover, the ability to conduct single subject statistics is additionally advantageous for both piloting experimental designs and for analyzing experiments which are limited by low subject numbers due to, amongst other things, the time required in preprocessing and by-hand analysis (e.g., 7T laminar/columnar studies). Importantly, we show that we can carry out single subject analysis with a relatively parsimonious experimental design, which does not require a large number of trials.

#### General considerations on FWER and power analysis

Though tMVPA was underpowered in simulations where 65% or fewer data points contained the effect of interest for both the group and single subject analyses, we argue that this is a potential strength rather than a weakness of our approach. While more likely to incur Type II errors (i.e. failing to reject H0), we would question the sensitivity, validity, and especially the generalizability of a method reporting statistical significance when only 65% or fewer data points display the effect being claimed. This argument becomes even more relevant in light of the recent emphasis of the scientific community on producing highly replicable studies, following the so called “replication crisis” (Schooler, 2014). We advocate the use of relatively more conservative statistical approaches, as we believe that overpowered statistical approaches can be regarded as one of the causes of the aforementioned replication crisis (Anderson & Maxwell, 2017). Furthermore, it is worth noting that the values estimated here (and the considerations that follow) are specific to our experimental settings and image acquisition parameters. We chose a stimulation paradigm (i.e. 850 ms visual stimulation; 4 trials per run) that is likely to lead to low evoked BOLD amplitude and, consequently, low experimental SNR (i.e. BOLD amplitude over trials measurement error). Under different stimulation regimes, such as longer stimulus presentation or block design experiments, we would expect higher statistical power or lower N to achieve the desired power. Moreover, at higher fields (i.e. 7T or above) the increase in both temporal and image SNR (Ugurbil, 2014) will be paired with a boost in statistical power. As such, the statistical power computed here in a relatively low SNR regime, represents a conservative estimate for the proposed approach.

#### Temporal multivariate approach to fMRI

Traditionally, due to the sluggish nature of the hemodynamic based BOLD signal (Boynton et al., 1996; S Ogawa et al., 1993), fMRI’s temporal resolution has traditionally been overlooked, deemed to be too inaccurate to measure the temporal dynamics of neural processing. More recently, however, a number of animal studies have begun exploring the temporal dimension of the BOLD signal. Functional images have been recorded in marmosets with a temporal resolution of 200 ms (Yen et al., 2018) and in rats with 40 ms (Silva & Koretsky, 2002). Furthermore, human recordings have suggested that increasing fMRI temporal resolution may reveal insights into the temporal dynamics of neural processing. For example, recent evidence put forward by Lewis et al (Lewis et al., 2016) suggest that fMRI can measure neural oscillatory activity at a much higher rate than previously suggested, specifically up to 1Hz. Accordingly, Siero et al. (Siero, J.C., Petridou, N., Hoogduin, H., Luijten, P.R., Ramsey, 2011) showed that, away from large draining, vessels the hemodynamic response function peaks ∼2 seconds earlier and is approximately 1 second narrower than previously reported, thus indicating that the neurovascular coupling occurs at a much shorted time-scale. Additionally, Vu et al.’s (Vu, A.T., Phillips, J.S., Kay, K., Phillips, M.E., Johnson, M.R., Shinkareva, S.V., Tubridy, S., Millin, R., Grossman, M., Gureckis, T., Bhattacharyya, R., Yacoub, 2016) work also advocates the importance of the BOLD temporal dimension. These authors showed that that, using MVPA, it is possible to extract timing information at fast TRs (i.e. 500 ms) that would otherwise be inaccessible (Vu, A.T., Phillips, J.S., Kay, K., Phillips, M.E., Johnson, M.R., Shinkareva, S.V., Tubridy, S., Millin, R., Grossman, M., Gureckis, T., Bhattacharyya, R., Yacoub, 2016).

These observations highlight the growing interest in the temporal dynamics of the BOLD signal, motivating the need for novel analytical tools specifically tailored to extract BOLD temporal information. Within this context, the method we developed is highly advantageous in that it incorporates the multivariate dimension in the temporal analysis of the BOLD signal, rendering potentially unexplored temporal features accessible. This mulitvariate dimension comes from considering the spatial pattern of BOLD activity across the voxels population within a given ROI at every time-point. As such, tMVPA extends the power of fMRI, which has historically been in the spatial domain, to the much less studied temporal dimension.

tMVPA thus allows investigating the temporal evolution of neural representation, which is incredibly valuable for exploring a wide range of phenomena, from visual illusions (Ernst & Bu, 2004; Schrater et al., 2004), real world scenes, and a variety of auditory paradigms (Baumann et al., 2010; Lee et al., 2012). As such, our method can be broadly applied to a large domain of stimulus paradigms.

Another interesting feature of tMVPA is the fact that paradigms utilizing active behavioral judgments of stimuli (as in Ramon et al. (Ramon et al., 2015)) may choose to align the analysis with either the stimulus onset or the behavioral response. This allows investigating response- as well as stimulus-locked modulations of neural representations over time.

It is also worth considering the nature of the effect being observed with tMVPA. Our technique measures multivariate activity at the population level accessible with fMRI [∼640,000 neurons (Lent et al., 2012)], and is as such constrained by the temporal lag of the BOLD signal (S Ogawa et al., 1993). While these constraints limit its temporal precision, especially relatively to the resolution available using invasive electrophysiological techniques (Meyers et al., 2015), tMVPA does provide valuable insights into the *relative* temporal dynamics of the neural processes captured with fMRI. In essence, while tMVPA won’t provide direct insights into the actual temporal window of neural processing, the careful investigation of temporal aspects of the BOLD signal could provide important information regarding the neural substrates of cognition (Seiji Ogawa et al., 2000; Smith et al., 2012). For example, the relative BOLD latency differences between experimental conditions can be related to diverse cognitive processes (Gentile et al., 2017; Henson et al., 2002).

tMVPA analysis already proved useful by revealing crucial differences in the temporal processing of familiar and unfamiliar faces in the left fusiform face area and in the bilateral amygdala (Ramon et al., 2015). Importantly, in Ramon et al. (Ramon et al., 2015) these differences would have remained undetected using traditional temporal univariate analysis techniques, as we did not observe significant differences between the average (across voxels and trials) BOLD time courses of familiar and unfamiliar faces. Accordingly, our simulations were carried out on synthetic data that were carefully generated with the absence of univariate amplitude differences across conditions (figure 3). We thus replicated what we originally showed in Ramon et al. (Ramon et al., 2015), namely, the ability of the tMVPA approach to detect genuine temporal multivariate effects or ones not driven by mere univariate amplitude differences.

It must be noted that the differences between this work and Ramon et al. (Ramon et al., 2015) are substantial both in terms of stimulation paradigm and MR acquisition parameters. Their functional scans were acquired using a repeated, single-shot echo planar imaging sequence with 3.5-mm isotropic voxel, a 64 × 64 matrix, a TE of 50 ms, TR of 1250 ms, FA of 90° and FOV of 224 mm. Moreover, Ramon et al. (Ramon et al., 2015) used a novel visual paradigm where a face stimulus was kept on screen for a duration of approximately 19 to 21 TRs, followed by a fixation period lasting 6 to 8 TRs. Yet, in spite of these differences, in both datasets our technique uncovered effects that were not detected when using traditional univariate methods focusing on amplitude differences between average time courses.

#### Validation on synthetic versus real data

It is important to consider that the multivariate data used to assess this technique were generated synthetically (see methods). Our technique was initially conceived for use with experimentally derived data (Ramon et al., 2015). As the goal of the present study is to assess the experimental parameters and conditions under which our technique is most useful, the ability to manipulate these variables is crucial and thus synthetic data is ultimately necessary. As previously mentioned, in an effort to generate a synthetic data set with realistic signal and noise properties, we used noise and signal estimates from real fMRI data. We approximated the fMRI signal by averaging BOLD time courses across voxels, trials, and conditions, and the amount of noise by measuring the variability (i.e. standard deviation) across voxels, trials and time-points. Hybrid approaches to synthetic data generations, such as the one implemented here, are highly beneficial (Welvaert & Rosseel, 2013). They provide full control over the data set, while preserving realistic signal to noise estimates and, according to (Welvaert & Rosseel, 2013), may represent the ideal data generation procedure for statistical validation. Our data generation approach, however, builds upon random sampling of variance and signal properties across voxels, ROIs, conditions, and subjects (see methods). This procedure effectively impairs the original temporal and spatial autocorrelation present in fMRI data. In the present study, we did not attempt to reinject temporal and spatial autocorrelation in the synthetic data. The reason behind this choice is twofold. Firstly, fMRI has multiple sources of noise (e.g. thermal, physiological, motion, task), each of which is characterized by different distributions and parameters, making it difficult to accurately and comprehensively model all noise sources. As such, an exhaustive model that allows generation of realistic fMRI noise has yet to be formulated. In order to introduce synthetic but *realistic* spatio-temporal auto-correlated noise in simulated fMRI data, there is first a need to formulate a comprehensive and realistic noise model. However, the quest for an exhaustive model for fMRI data (including noise) generation is challenging enough to require a study in and of itself tailored to tackle this specific endeavor (Davis et al., 2014) and, as such, is well beyond the scope of this article. Additionally, given the lack of a “ground-truth” noise model, noise estimates may be inaccurate or misrepresent the contribution of difference noise sources and, as such, noise injection may have a negative impact on the validation procedure as a whole. Secondly, we argue that the impact of spatio-temporal auto-correlated noise is minimal within these specific settings. The structure of the stRDMs when considering real, as opposed to synthetic, data can be seen in figure 1. Patches of similarity (cool colors) and dissimilarity (warm colors) exist in clusters of approximately 3-4 TRs. Such structure is due to the inherent spatiotemporal autocorrelation present in the BOLD signal, which is not dependent on experimental manipulations. Rather, it is a direct outcome of the HRF response properties. Specifically, BOLD activation for all voxels will synchronously rise for approximately the first 6 seconds after stimulus onset (varying depending on stimulus presentation time), and then decrease for the following 6 seconds, thus generating the structure visible in the matrices in figure 1. This structure will therefore be shared across conditions and subtracted out when performing the linear contrast between the stRDMs across conditions (see methods). As such, the inherent presence of autocorrelation in fMRI data, which is shared across conditions, becomes irrelevant in evaluating the validity of our validation procedure.

## Conclusion

In summary, we have developed a method for examining the representational content of fMRI data as a function of time, whereby enabling the investigation of the temporal evolution of neural representation. The method, that builds upon fMRI most recognized strength – namely its spatial resolution – to analyze BOLD temporal dynamics, consists of creating Single Trial Representational Dissimilarity Matrices (stRDMs) to measure the dissimilarity between the neural representations elicited by each acquired time point of a BOLD time course. We also introduced an expanding, sliding window method for inferring statistical significance. We validated our temporal multivariate pattern analysis (tMVPA) in both group and single subject settings using synthetically generated data. Our results show that we achieve adequate power FWER in both contexts. Along with the addition of a multivariate dimension to BOLD temporal analyses, tMVPA permits performing single subject’s inferential statistics by considering single trial distributions. Importantly, single subject analysis can be reliably implemented with a parsimonious experimental design that requires as little as 12 trials per condition across all runs. Furthermore, we show that, both in simulated as well as real settings (see Ramon et al. (Ramon et al., 2015)), our tMVPA is capable of detecting multivariate effects between experimental conditions in the absence of univariate amplitude differences. The technique presented here expands on traditional multivariate fMRI analyses, facilitating investigations of the evolution of neural representations over time.

## Footnotes

1 Note that correlational distance 1-r can be conceptualized as distance between 2 points in a multidimensional space. In the same vein, we can think of increase in correlation (and therefore decrease in correlational distance) between these 2 points as a rotation of axis of the the multidimensional space for point 1

